# The inactivation of enzymes belonging to the central carbon metabolism, a novel mechanism of developing antibiotic resistance

**DOI:** 10.1101/823013

**Authors:** Teresa Gil-Gil, Fernando Corona, José Luis Martínez, Alejandra Bernardini

## Abstract

Fosfomycin is a bactericidal antibiotic, analogous to phosphoenolpyruvate (PEP) that exerts its activity by inhibiting the activity of MurA. This enzyme catalyzes the first step of peptidoglycan biosynthesis, the transfer of enolpyruvate from PEP to uridine-diphosphate-N-acetylglucosamine. Fosfomycin is increasingly used in the last years, mainly for treating infections caused by Gram-negative multidrug resistant bacteria as *Stenotrophomonas maltophilia*, an opportunistic pathogen characterized by its low susceptibility to antibiotics of common use. The mechanisms of mutational resistance to fosfomycin in *S. maltophilia* were studied in the current work. None of the mechanisms so far described for other organisms, which include the production of fosfomycin inactivating enzymes, target modification, induction of alternative peptidoglycan biosynthesis pathway and the impaired entrance of the antibiotic, are involved in the acquisition of such resistance by this bacterial species. Rather the unique cause of resistance in the studied mutants is the mutational inactivation of different enzymes belonging to the Embden-Meyerhof-Parnas central metabolism pathway. The amount of intracellular fosfomycin accumulation did not change in any of these mutants showing that neither the inactivation nor the transport of the antibiotic were involved. Transcriptomic analysis also showed that the mutants did not present changes in the expression level of putative alternative peptidoglycan biosynthesis pathway genes neither any related enzyme. Finally, the mutants did not present an increased PEP concentration that might compete with fosfomycin for its binding to MurA. Based on these results, we describe a completely novel mechanism of antibiotic resistance based on the remodeling of *S. maltophilia* metabolism.

**Significance:** Antibiotic resistance (AR) has been largely considered as a specific bacterial response to an antibiotic challenge. Indeed, its study has been mainly concentrated in mechanisms that affect the antibiotics (mutations in transporters, the activity of efflux pumps and antibiotic modifying enzymes) or their targets (i.e.: target mutations, protection or bypass). Usually, AR-associated metabolic changes were considered to be a consequence (fitness costs) and not a cause of AR. Herein, we show that strong alterations in the bacterial metabolism can also be the cause of AR. In the study here presented, *Stenotrophomonas maltophilia* acquires fosfomycin resistance through the inactivation of glycolytic enzymes belonging to the Embden-Meyerhof-Parnas. Besides resistance to fosfomycin, this inactivation also impairs the bacterial gluconeogenic pathway. Together with previous work showing that AR can be under metabolic control, our results provide evidence that AR is intertwined with the bacterial metabolism.

## Introduction

Antibiotic resistance can be considered as a chemical problem. To be active, an antibiotic requires to reach its target at concentrations high enough for inhibiting its activity. Any process or situation that either reduces the effective concentration of the antibiotic or the antibiotic-target affinity should lead to antibiotic resistance. In agreement with this situation, classical, so far described, mechanisms of resistance (1) include elements that diminish the antibiotic concentration like efflux pumps (2), antibiotic inactivating enzymes (3) or changes in the antibiotic transporters (4). Concerning the target, elements that reduce its affinity with the antibiotic include mutations (5), target protection (6), bypass (7) or replacement (8) and eventually increased target expression (9). Studies on intrinsic resistome have shown that, in addition to these classical resistance determinants, the susceptibility to antibiotics of a bacterial species depends on the activity of several elements encompassing all functional categories (10-12). However, little is still known about the interplay between bacterial metabolism and the acquisition of antibiotic resistance (13). In the current article, we explore this feature analyzing *S. maltophilia* fosfomycin resistant mutants. Fosfomycin is a phosphonic acid derivative that contains an epoxide and a propyl group, chemically analogous to phosphoenolpyruvate (PEP), which explains its mechanism of action (14). The enzyme MurA (UDP-*N*-acetylglucosamine enolpyruvyl transferase), which catalyzes the first step in peptidoglycan biosynthesis (15), the transfer of enolpyruvate from PEP to uridine diphospho-N-acetylglucosamine, is the only known fosfomycin target. Fosfomycin binds covalently to a cysteine residue in the active site of MurA, which renders MurA inactivation. As a consequence of MurA inactivation, the peptidoglycan precursor monomers accumulate inside the cell, peptidoglycan cannot be synthesized and this leads to bacterial cell lysis and death (16).

Different molecular mechanisms leading to fosfomycin resistance have been identified (17). Some of them impairing the fosfomycin/MurA interaction. Some allelic variants of MurA found in pathogens intrinsically resistant to fosfomycin such as *Mycobacterium tuberculosis, Borrelia burgdorferi* or *Chlamydia* sp. (15, 18-20) do not contain a cysteine in their active site, and therefore they are not fully inhibited by fosfomycin. In the case of organisms containing a fosfomycin-sensitive MurA allele, mutations in *murA* can be selected (15, 21, 22) and increased synthesis of MurA also confers a resistance phenotype (23, 24). Also, the presence of an alternative route of peptidoglycan synthesis, as it happens in *Pseudomonas putida* and *P. aeruginosa*, may allow circumvent the activity of fosfomycin by recycling the peptidoglycan without the need of *de novo* synthesis by the enzyme MurA (7). Concerning mechanism involving a reduction in the intracellular concentration of the antibiotic, resistance can be achieved as the consequence of changes in the entrance of fosfomycin inside bacterial cell. The main cause of this impaired uptake is the selection of mutations in any of the genes encoding the sugar phosphate transporters GlpT and UhpT, which are the gates for fosfomycin entrance (25, 26). To note here that expression of these transporters is under metabolic control, in such a way that situations where the nutritional bacterial status favors the use of sugar phosphates (as intracellular growth) increase fosfomycin activity (27, 28). Finally, in other cases, fosfomycin is inactivated by fosfomycin modifying enzymes as FosA, FosB and FosX. (29-32). All the already known mechanisms of fosfomycin resistance fit in the classical categories of resistance elements (see above). However, the results presented in the current article support that none of them are involved in the acquisition of resistance by *S. maltophilia.* In this bacterial species, fosfomycin resistance was acquired due to mutations in genes encoding enzymes of the Embden-Meyerhof-Parnas (EMP) metabolic pathway. It has been suggested that antibiotic resistance can be inter-linked to bacterial metabolism (33, 34). However, with very few exceptions (35), the mutational inactivation of genes encoding enzymes of the central carbon metabolism has not been considered to be a significant cause of antibiotic resistance in bacterial pathogens (34, 35). Our article hence shed light in the crosstalk between antibiotic resistance and central carbon metabolism in *S. maltophilia*.

## Results

### Selection of *S. maltophilia* fosfomycin resistant mutants and identification of the mutations involved

In order to isolate single-step *S. maltophilia* fosfomycin resistant mutants, around 10^8^ bacterial cells were seeded on selection plates containing fosfomycin (1024 µg/ml). Four single-step fosfomycin resistance mutants, hereafter dubbed FOS1, FOS4, FOS7 and FOS8, were selected for further studies. The MIC of the mutants to fosfomycin was determined. In all cases, the MICs of fosfomycin were higher in the mutants than in the wild-type strain (Table 1).

**Table 1.** MICs (µg/ml) of fosfomycin for the resistant mutants and their corresponding complemented strains

| Strain | Plasmid used for complementation |  |  |  |  |  |
| --- | --- | --- | --- | --- | --- | --- |
|  | None | pSEVA234 | pSEVA234<br>*eno* | pSEVA234<br>*gpmA* | pSEVA234<br>*gapA* | pSEVA234<br>*pgk* |
| **D457** | 192 | 192 | 128 | 128 | 128 | 192 |
| **FOS1** | >1024 | >1024 | 256 | ND | ND | ND |
| **FOS4** | >1024 | >1024 | ND | 256 | ND | ND |
| **FOS7** | >1024 | >1024 | ND | ND | 128 | ND |
| **FOS8** | >1024 | >1024 | ND | ND | ND | 192 |

ND: Not done, each strain was complemented with the wild-type allele of the corresponding mutated gene.

The genomes of these mutants were fully sequenced and compared with that of the parental wild-type strain D457. Five different mutations were detected. FOS4, FOS7 and FOS8 carried one mutation, while FOS1 harbored two mutations. One of them (in *rne*, SMD_RS14705:c.G1464T:p.E488D) was discarded because it was predicted to be neutral using the Provean predictor (0.41 score). Notably, each mutant contains a different mutation, but all four were found in genes encoding enzymes of EMP metabolic pathway, namely *eno, gpmA, gapA* and *pgk* (Table 2). To further confirm the presence of each of these mutations in the mutant strains, the corresponding genomic regions were amplified by PCR and the amplicons were Sanger-sequenced. The presence of the mutations was confirmed in all cases.

**Table 2.** SNPs mapped in the mutants presenting low susceptibility to fosfomycin.

| Mutant | Position of the SNP<br>reference sequence | SNP | Locus | Gene | Amino acid<br>change | Product | Old locus<br>tag | Provean<br>Score* |
| --- | --- | --- | --- | --- | --- | --- | --- | --- |
| **FOS1** | 1.829.461 | A:T>G:C | SMD_RS08765 | *eno* | p.D398G | Enolase | SMD_1655 | -6.77 |
| **FOS4** | 1.411.119 | C:G>T:A | SMD_RS06650 | *gpmA* | p.P212L | Phosphoglycerate mutase | SMD_1268 | -9.88 |
| **FOS7** | 3.798.418 | G:C>C:G | SMD_RS17680 | *gapA* | p.D296G | Glyceraldehyde-3-phosphate<br>dehydrogenase | SMD_3406 | -3.37 |
| **FOS8** | 3.793.349 | G:C>A:T | SMD_RS17665 | *pgk* | p.Q50ST | Phosphoglycerate kinase | SMD_3403 |  |

\*Provean deleterious score threshold of the changes is -2.5.
ST: Stop mutation

Although no other mutations seemed to be the cause of the resistance of the studied mutants, the wild-type allele of the corresponding mutated gene was introduced in each mutant strain to get a functional validation of the effect of these mutations in the susceptibility to fosfomycin of *S. maltophilia*. As shown in Table 1, introduction of the wild-type forms of such genes fully restores the susceptibility of the analyzed *S. maltophilia* fosfomycin resistant mutants to the level of the wild-type strain. These results indicate that the fosfomycin resistance of these mutants is solely due to the mutation of genes encoding enzymes of the EMP metabolic pathway. In addition, susceptibility to other antibiotics was tested in the fosfomycin resistant mutants. No significant changes between the wild-type strain and the mutants were observed for any of the tested antibiotics (Table S1), which strongly suggests that these mutations in genes coding enzymes of the central metabolism are fosfomycin-specific resistance mutations.

### Model of *S. maltophilia* central metabolism

As a first step for deciphering how the mutations in genes encoding enzymes of the EMP metabolic pathway may impact *S. maltophilia* physiology, a metabolic map of the central metabolism, which generate energy and precursors to form biomass (36), was modeled for *S. maltophilia*. The EMP pathway is the best analyzed glycolytic route. It is based on the sequential activity of ten individual enzymes. The first five form the upper glycolysis (Glk, Pgi, Pfk, Alf1, TpiA) in which, using ATP, hexoses are converted into trioses phosphate; whereas in the lower glycolysis (GapA, Pgk, GpmA, Eno, PykA), pyruvate is formed from the trioses phosphate, at the same time that NADH and ATP are generated. The pyruvate obtained is decarboxylated by the action of pyruvate dehydrogenase complex and enters as acetyl-CoA to the tricarboxylic acids (TCA) cycle (37). The EMP pathway may also function in a gluconeogenic regime, forming hexoses phosphate from trioses phosphate (38). All enzymes of the EMP pathway were identified in *S. maltophilia* D457 (Figure 1 and Figure S1). Moreover, the Entner-Doudoroff (ED) route, another glycolytic pathway that also forms trioses phosphate from hexoses phosphate, is present as well in *S. maltophilia*. It is important to notice that two enzymes of the central metabolism of D457, GpmA and Eno, present isoenzymes capable of carrying out the same chemical reaction. As shown in Figure 1, all the fosfomycin resistance mutations are located in genes encoding enzymes of the lower glycolytic pathway.

**Figure 1.**
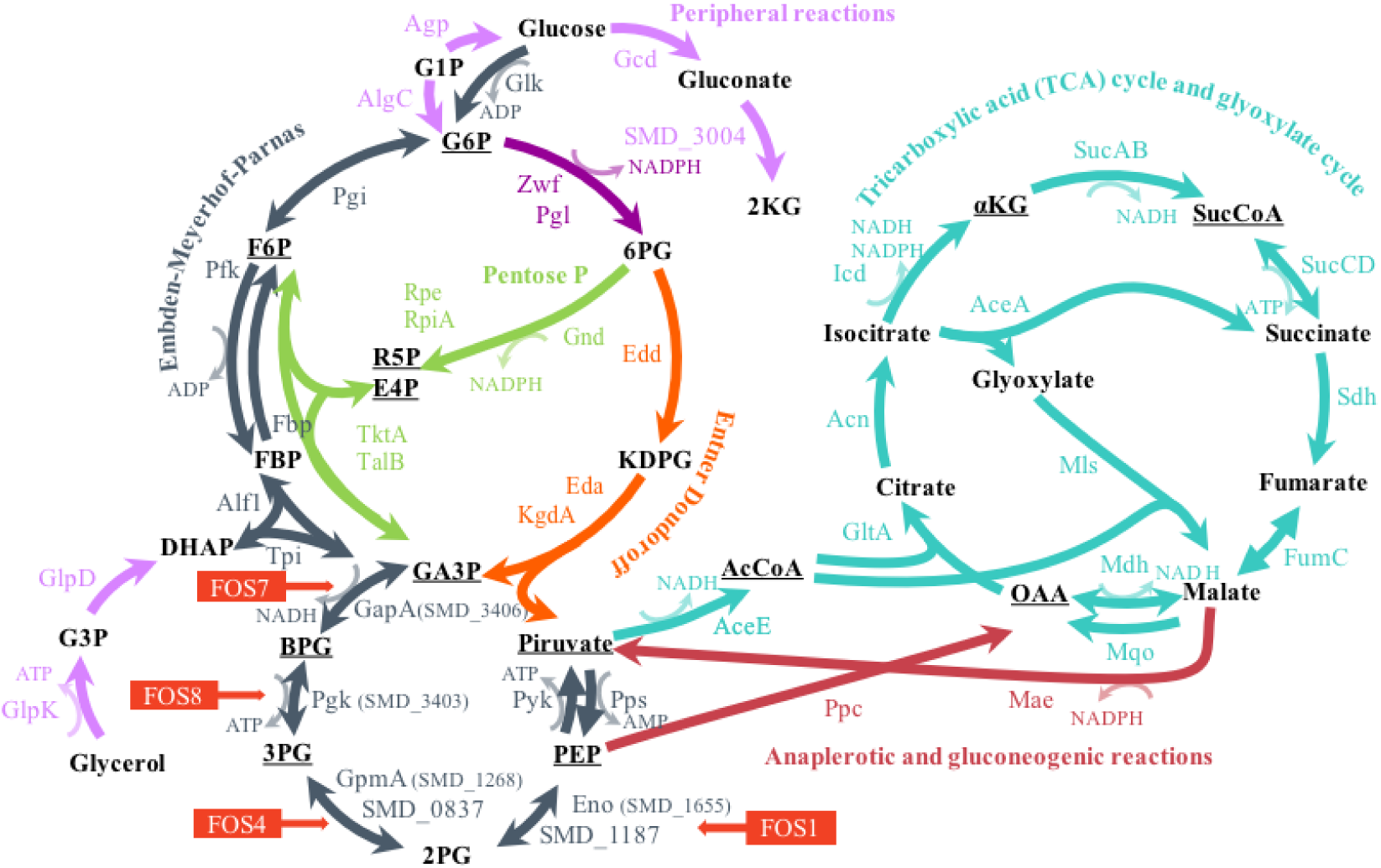
Central metabolism of *S. maltophilia* D457. Schematic representation of the main pathways of the central metabolism: glycolysis (Entner-Doudoroff and Embden-Meyerhof-Parnas), tricarboxylic acid cycle and glyoxylate cycle; pentose phosphate pathway; and anaplerotic and gluconeogenic reactions, as well as peripheral reactions. Underlined are the essential precursors for the biomass formation (36). The mutated enzyme in each FOS mutant is indicated with the name of the corresponding mutant. The name of substrates and products, and the code of the enzymes is shown in Supplemental Material, Figure S1.

### Fosfomycin resistance mutations impair the activity of enzymes of the *S. maltophilia* central metabolism

To determine whether or not the mutations cause a loss of function of the encoded proteins, the enzymatic activity of Gap, Pgk, Gpm and Eno was measured in the mutants and in the wild-type strain. As shown in Figure 2, Gap activity decreased by 93 % in the FOS7 mutant, Pgk activity decreased by 100 % in FOS8, Gpm activity decreased by 65% in FOS4 and Eno activity decreased by 100 % in FOS1, in relation to the parental strain. Thus, every mutation causes a loss of function of the encoded gene.

**Figure 2.**
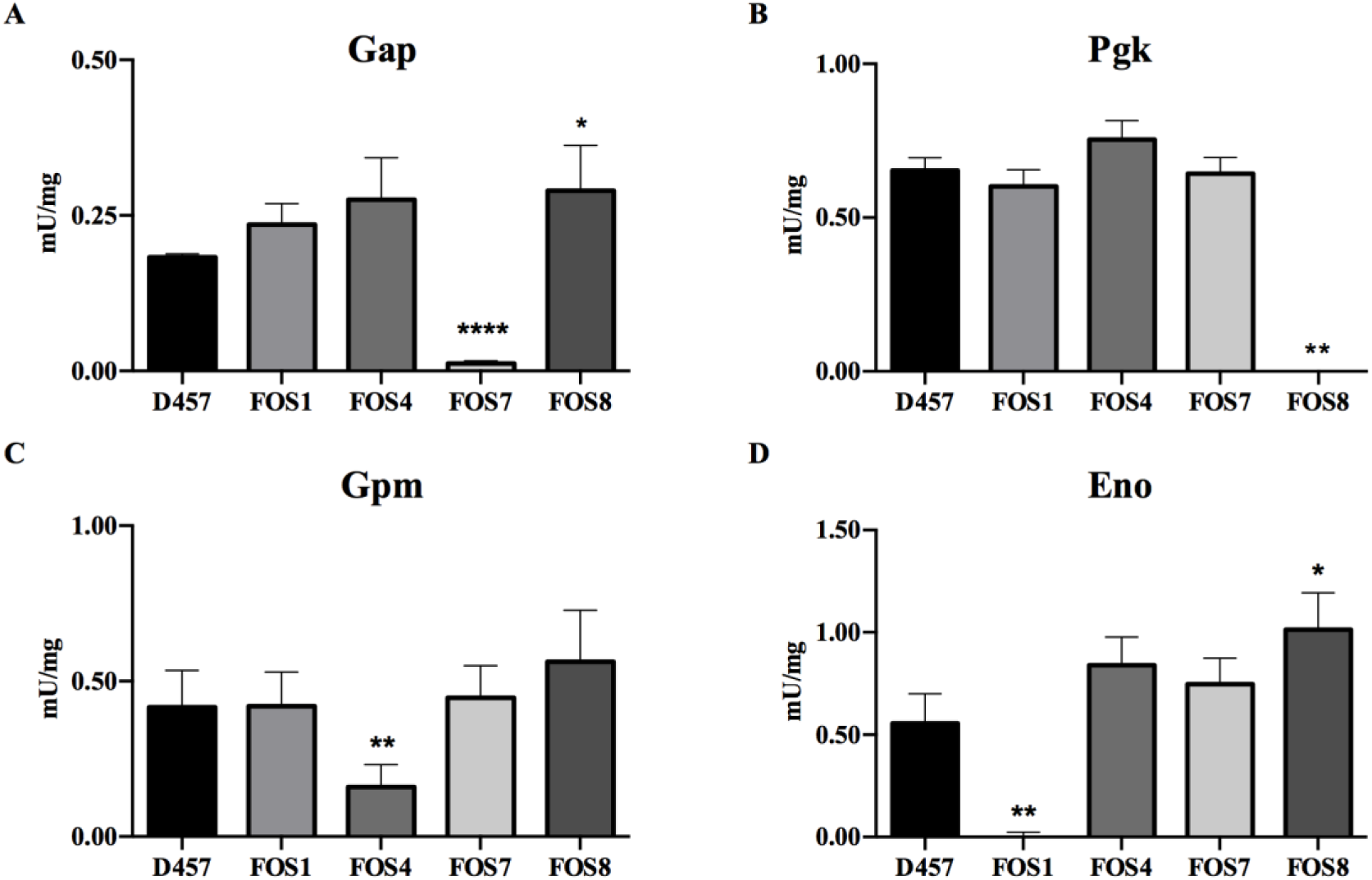
Enzymatic activity of the lower glycolysis enzymes in the D457 parental strain and in the fosfomycin resistant mutants. **A) Gap** Glyceraldehyde-3P dehydrogenase activity. **B) Pgk** Phosphoglycerate kinase activity. **C) Gpm** Phosphoglycerate mutase activity. **D) Eno** Enolase activity. Error bars indicate standard deviations of the results from three independent replicates. **\*** Indicates *P* < 0.02, ** *P* < 0.002 and **\*\*\***\* *P* < 0.0001 calculated by unpaired two tail t-test. As shown, each of the mutants present an impaired activity of the enzyme encoded by the mutated gene.

To elucidate if the reduced activity of these enzymes in the mutants may produce a relevant metabolic shift in *S. maltophilia*, the activities of the main dehydrogenases of the central metabolism of *S. maltophilia* D457, which are indicative of the general physiological state of the cell, including its redox balance, were measured. In particular, the activities of the glucose-6P dehydrogenase (Zwf), which connects the glucose-6P with the ED and pentose phosphate (PP) pathways, and isocitrate dehydrogenases (Icd NAD^+^ and Icd NADP^+^) activity, from the TCA cycle, were determined. The activity of the enzyme Zwf increased by 1.5 to 2.5-fold in the four fosfomycin resistant mutants (Figure 3) as compared with the wild-type D457 strain, whereas the activities of either Icd NAD^+^ or Icd NADP^+^ did not change in any of the studied mutants.

**Figure 3.**
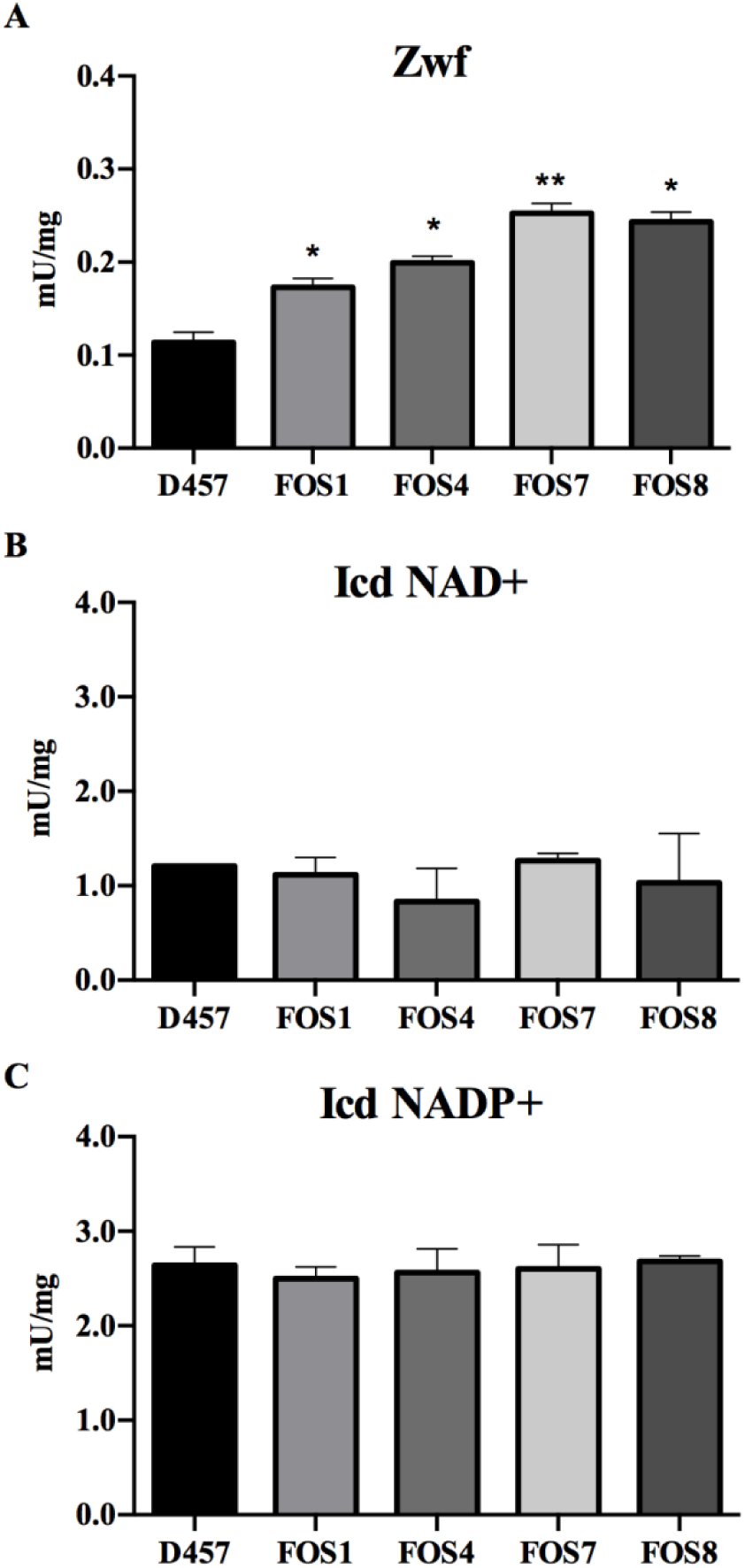
Enzymatic activity of dehydrogenases from *S. maltophilia* central metabolism of D457 parental strain and fosfomycin resistant mutants. **A) Zwf** Glucose-6P dehydrogenase activity. **B) Icd (NAD**^**+**^) Isocitrate dehydrogenase NAD^+^ activity. **C) Icd (NADP**^**+**^) Isocitrate dehydrogenase NADP^+^ activity. Error bars indicate standard deviations of the results from three independent replicates. As shown, the activity of Zwf is higher in the fosfomycin resistant mutants. * Indicates *P* < 0.02 and ** *P* < 0.005 calculated by unpaired two tail t-test.

### Fosfomycin resistance is not the consequence of a metabolic rearrangement that modifies *S. maltophilia* susceptibility to oxidative stress

It has been proposed that the activity of antibiotics may depend on the bacterial oxidative response (39). One of the key elements in such response is Zwf, an enzyme with a critical role in the supply of NADPH, which is a relevant cofactor for maintaining cellular redox balance (40, 41). We have observed that this enzyme presented an increased activity in the mutants as compared with the wild-type strain (see above). To address if this increased activity might be the reason for fosfomycin resistance, *zwf* was inactivated in the FOS4 and FOS7 mutants and in the D457 wild-type strain. The inactivation of *zwf* causes a slight increase in MIC levels of fosfomycin from 192 to 256 μg/ml in D457 wild-type strain, whereas this inactivation does not change fosfomycin susceptibility in the tested mutants.

Besides, the role of the mutations in the response to oxidative stresses was tested by analyzing the susceptibility of the mutants to H_2_O_2_ and menadione. As shown in Table 3, mutations conferring fosfomycin resistance did not alter the susceptibility of *S. maltophilia* to these compounds, whereas, as expected, *zwf* inactivation causes an increase in the susceptibility to these oxidative stressors. These results indicate that the susceptibility to fosfomycin of *S. maltophilia* mutants with defective lower glycolysis enzymes is a specific phenotype, not due to a change in the oxidative stress response.

**Table 3.** Susceptibility to oxidative stress of the analyzed mutants.

| Strains | H<sub>2</sub>O<sub>2</sub> | Menadione |
| --- | --- | --- |
| **D457** | 4.4 ± 0.2 | 1.9 ± 0.2 |
| **FOS1** | 4.5 ± 0.3 | 1.9 ± 0.3 |
| **FOS4** | 4.0 ± 0.3 | 2.1 ± 0.3 |
| **FOS7** | 3.9 ± 0.4 | 1.5 ± 0.5 |
| **FOS8** | 4.1 ± 0.1 | 2.0 ± 0.8 |
| **D457 $\Delta zwf$** | 5.6 ± 0.8 | 2.6 ± 0.3 |
| **FOS4 $\Delta zwf$** | 5.0 ± 0.5 | 4.9 ± 0.2 |
| **FOS7 $\Delta zwf$** | 6.6 ± 0.7 | 4.7 ± 0.6 |

### The impaired activity of EMP enzymes is associated with *S. maltophilia* antibiotic resistance

Our results strongly suggest that the cause of fosfomycin resistance in the studied mutants is a reduced activity of the enzymes of the lower glycolysis pathway in *S. maltophilia*. However, it is still possible that these enzymes may present moonlighting activities in this bacterial species besides its metabolic role, which could be associated with the antibiotic resistance in a metabolic independent manner (42, 43). This possibility is supported by the fact that, while mutations in these genes are easily selected in *S. maltophilia*, the information present in the Profiling of the *E. coli* Chromosome (PEC) database, the Keio library and the Transposon-directed insertion site sequencing (TraDIS) database (44-46) support that they are highly relevant (eventually essential) in *E. coli*.

To determine if the recovery of the glycolytic activity, independently of a putative additional activity of the *S. maltophilia* inactivated enzymes, could be on the basis of the observed antibiotic resistance phenotype, a partial version of the Glucobrick II, containing the *E. coli* genes *gapA, pgk, gpmA* and *eno* was introduced in the *S. maltophilia* fosfomycin resistant mutants and in the wild-type strain and the susceptibility to fosfomycin of these strains was measured. By this approach the enzymatic activity, here provided by the *E. coli* orthologues of the *S. maltophilia* inactivated genes, was decoupled from another potential activity of such *S. maltophilia* proteins. As shown in Figure 4, the expression of the *E. coli* GapA-Pgk-GpmA-Eno enzymes increased the susceptibility to fosfomycin of all FOS mutants, although the levels achieved were not the same as those of the wild-type strain. This partial complementation of the phenotype of resistance strongly supports that the absence of enzymatic activity of the analyzed EMP enzymes contribute to fosfomycin resistance in *S. maltophilia*.

**Figure 4.**
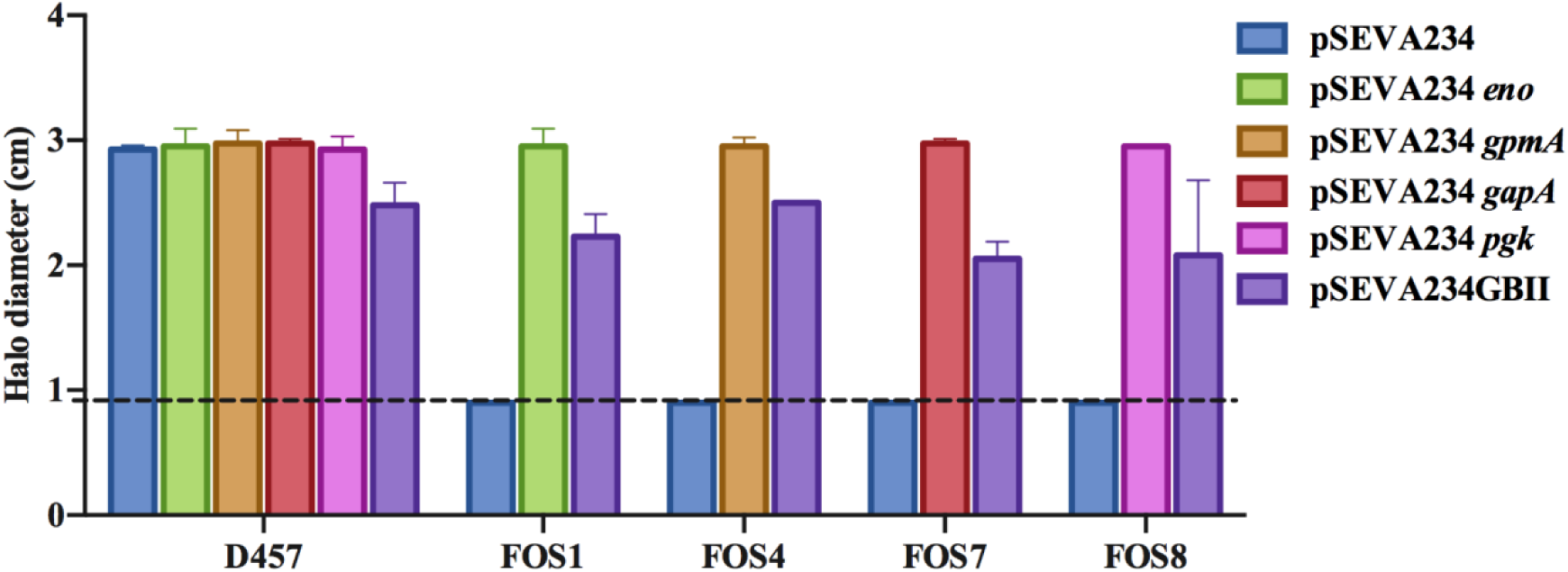
Fosfomycin susceptibility of the fosfomycin resistant mutants complemented with either *S. maltophilia* D457 or *E. coli* K-12 enzymes. The halo inhibition diameter of fosfomycin disks is shown for the wild-type D457 strain and the four mutants. In all cases, the results are shown for the strains containing either the pSEVA234 backbone used for cloning, the corresponding wild-type alleles of *S. maltophilia* genes (*eno, gpmA, gapA, pgk*) or a partial version of Glucobrick II (GBII) with *E. coli* genes *gapA, pgk, gpmA* and *eno.* Dashed line at 0.9 cm indicates the diameter of the disk. Error bars indicate standard deviations of the results from three independent replicates.

### Fosfomycin resistance of mutants defective in EMP enzymes is not the consequence of an increased production of phosphoenolpyruvate

Fosfomycin inhibits the action of MurA because it is structurally similar to PEP, one of the substrates of this enzyme. The EMP enzymes associated with fosfomycin resistance that are inactivated in the *S. maltophilia* fosfomycin resistant mutants present reversible activity and belong to a pathway that leads to either PEP biosynthesis or consumption depending on the metabolic regime. It might be then possible that the inactivation of such enzymes may change the intracellular PEP concentrations, affecting the binding of fosfomycin to the active site of MurA through a possible competition between PEP and fosfomycin, which may render a reduced susceptibility to fosfomycin. To analyze this possibility, the concentration of PEP was analyzed in the wild-type D457 strain and in the fosfomycin resistant mutants. In none of the mutants an increase in the intracellular concentration of PEP was observed, ruling out the hypothesis that the cause of the reduced susceptibility to fosfomycin of the analyzed mutants is an increased production of PEP.

### Fosfomycin resistance mutations impair the gluconeogenic pathway of *S. maltophilia*

The mutations selected in presence of fosfomycin compromise the activity of relevant enzymes of *S. maltophilia* central metabolism. It is then expected, this would have relevant physiological consequences. To have a general scope of these consequences, the growth of *S. maltophilia* mutants and of wild-type parental strain under different conditions was measured. Just small differences in growth among the tested strains were observed for bacteria growing in rich LB medium (Figure 5A), indicating these mutations do not impose a relevant general, non-specific, fitness cost. In addition, the mutants can grow using glucose, which imposes a glycolytic metabolism, although in the case of FOS1 and FOS8 at a different rate (Figure 5B). Nevertheless, the mutants were unable to grow using succinate as the carbon source (Figure 5C). This impaired growth in succinate was not observed when the mutants were complemented with either the wild-type allele of each of the mutated enzymes or the *E. coli*-derived Glucobrick II (Figure S2). The blocking of any of the enzymes of the EMP pathway, between triose phosphate isomerase and pyruvate kinase, breaks the amphibolic process in two branches. These branches work in opposite directions, starting either from glucose or from pyruvate to provide energy or biosynthetic intermediates (47). Since *S. maltophilia* additionally displays the one-direction ED pathway for glucose catabolism, fosfomycin low susceptibility mutants can grow in minimal medium with glucose. Nevertheless, succinate as exclusive carbon source does not support growth of the mutants because gluconeogenesis and consequently synthesis of hexose phosphates are impaired.

**Figure 5.**
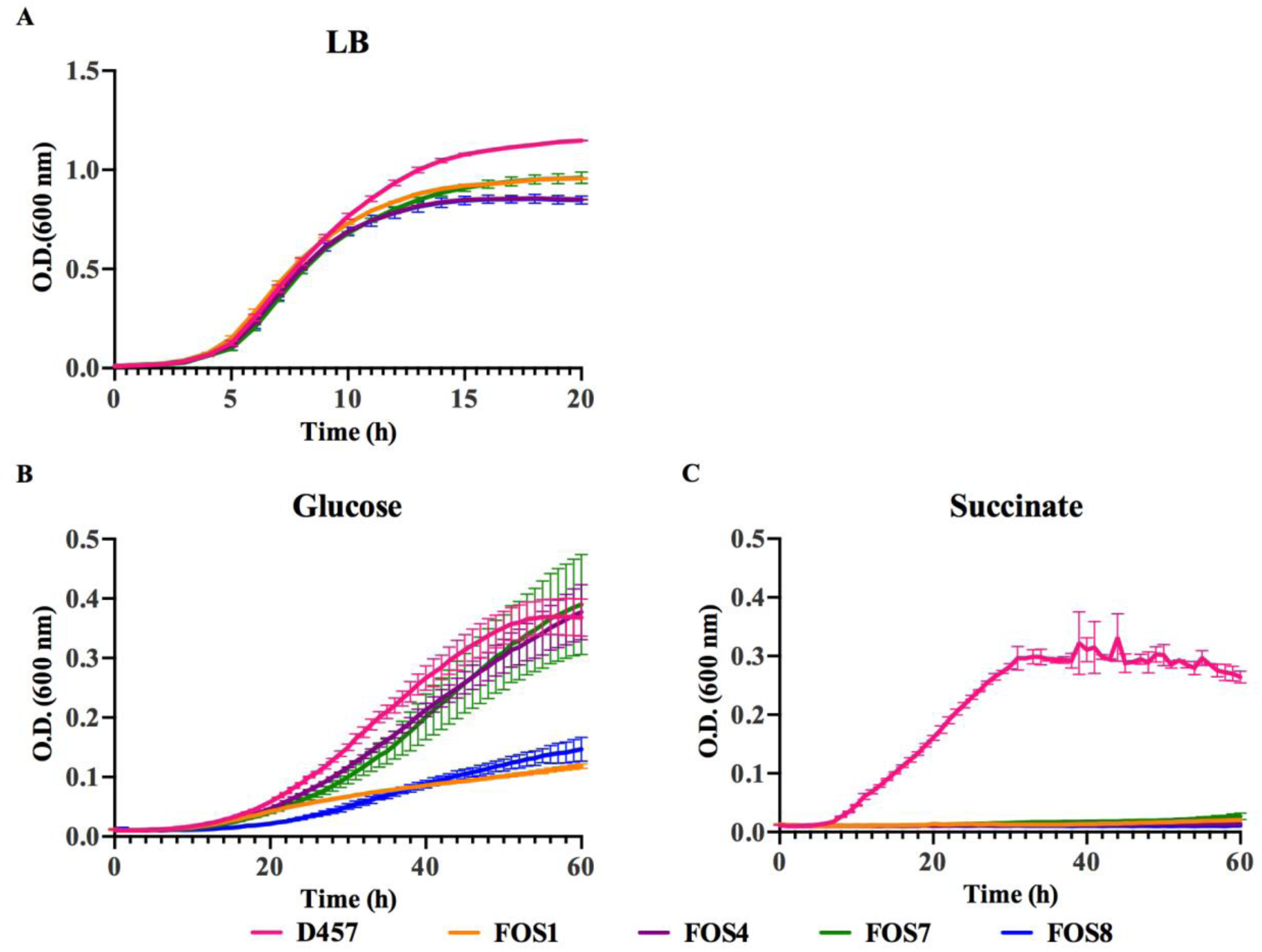
Effect of fosfomycin resistance mutations in the growth of *S. maltophilia* in either LB, glucose or succinate. **A)** Growth of the different strains in LB. **B)** Growth in SMMM containing glucose 40 mM. **C)** Growth in SMMM containing succinate 40 mM. As shown, the fosfomycin resistant mutants are strongly impaired for growing in succinate, whereas the impairment for growing in glucose, particularly of FOS4 and FOS7, was lower. The effect of the mutations in the growth in LB is limited. Error bars indicate standard deviations of the results from three independent replicates.

### Fosfomycin resistant mutants do not present an altered intracellular accumulation of fosfomycin

While the primary cause of fosfomycin resistance in *S. maltophilia* is the inactivation of EMP enzymes, it might be possible that such inactivation impairs the accumulation of the antibiotic within the cell, which could be due to either a reduced uptake or to the degradation of the antibiotic. A search of possible fosfomycin transporters in *S. maltophilia* D457 was carried out using Blast (48) with the sequences of fosfomycin transporters UhpT and GlpT. This search did not identify any possible transporter of hexose and triose phosphates in the genome of *S. maltophilia* D457, orthologous to those known in other microorganisms. Nevertheless, these sugars can still be internalized by alternative transporters. To address this possibility, the different strains were grown in SMMM containing either glucose-6P or glycerol-3P as sole carbon sources. Despite *S. maltophilia* harbors the orthologues of the enzymes required for the catabolism of glucose-6P and glycerol-3P, none of the strains were able to grow using these sugars as unique carbon sources, conditions at which *E. coli* can grow (Figure S3). This result suggests that *S. maltophilia* lacks glucose-6P and glycerol-3P transporters, which are the regular gates for fosfomycin entrance in other pathogens. Besides, the search of fosfomycin modifying enzymes in *S. maltophilia* D457 genome did not allow to detect any gene homologous to the fosfomycin resistance proteins (FosA, FosB, FosX, FomA, FomB and FosC) so far described in the literature.

Despite *S. maltophilia* genome does not harbor genes encoding neither the canonical fosfomycin transporters nor already known fosfomycin inactivating enzymes, it might still be possible that other (still unknown) elements may contribute to an impaired accumulation of the antibiotic inside the mutants. To analyze this possibility, the intracellular accumulation of fosfomycin in the different strains was measured (49) after one hour of incubation with 2 mg/ml fosfomycin in exponential growth phase cultures. As a control, *E. coli* K-12 and a deletion mutant on the fosfomycin transporter UhpT (44), as well as *P. aeruginosa* PA14 and insertion mutants on the fosfomycin transporter GlpT or the fosfomycin resistance protein FosA (50) were used. As shown in Figure 6, the amount of intracellular fosfomycin is lower both in *E. coli* and *P. aeruginosa* when their respective fosfomycin transporters (GlpT and UhpT) are inactivated. Conversely, an increased fosfomycin concentration was observed in the FosA mutant relative to the parental PA14 strain, which supports the validity of these assays. Nevertheless, the intracellular concentrations of fosfomycin were similar in the *S. maltophilia* D457 wild-type strain and in the isogenic fosfomycin resistant mutants. These results suggest that the resistance to fosfomycin of the FOS mutants is not due to a reduced intracellular concentration of this antibiotic. Notably, fosfomycin accumulation in *S. maltophilia* is much lower than that found in *E. coli* or *P. aeruginosa*. Indeed, intracellular fosfomycin concentration in *S. maltophilia* is in the range of that observed for the GlpT-defective *P. aeruginosa* mutant. This low intracellular concentration, likely associated to the lack of canonical antibiotic transporters, could be the cause of the intrinsic lower susceptibility of *S. maltophilia* D457 to fosfomycin compared to *E. coli* K-12 and *P. aeruginosa* PA14 (49, 51, 52).

**Figure 6.**
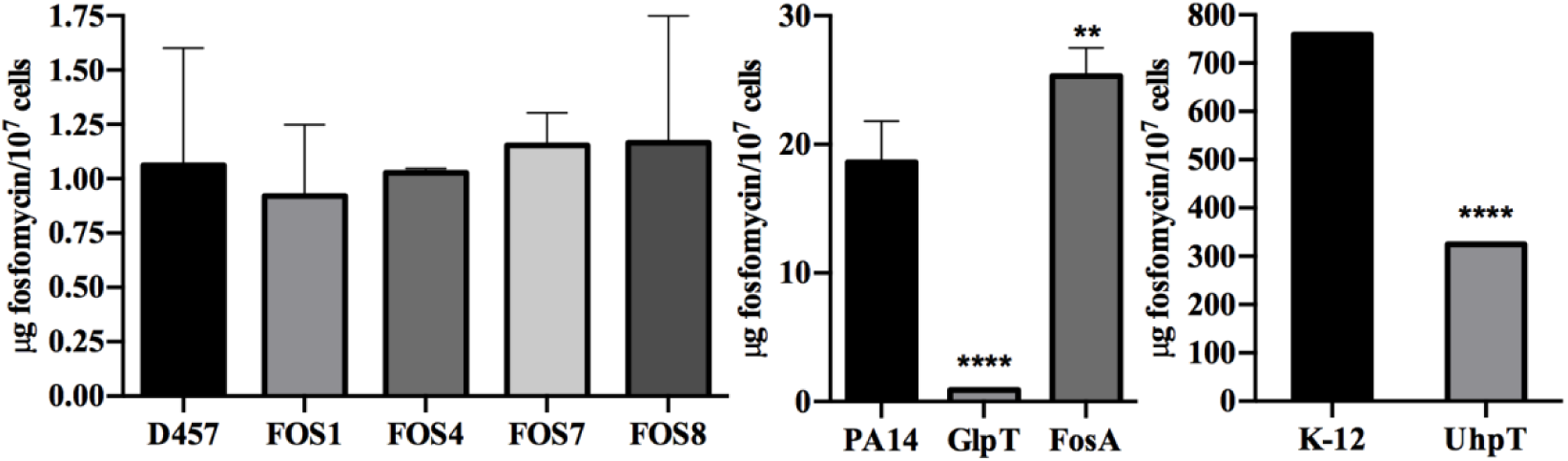
The intracellular concentration of fosfomycin does not change in the fosfomycin resistant mutants. Comparison of the fosfomycin intracellular concentration between the mutants and the parental strain. There is not a deficiency in the fosfomycin transport in the mutants that determines its resistance or a fosfomycin modifying enzyme involved in this resistance. PA14 and its mutant GlpT and FosA and *E. coli* K-12 and its mutant UhpT were used as controls of the assay. Error bars indicate standard deviations of the results from three independent replicates. ** Indicates *P* < 0.001 and **** *P* < 0.0001 calculated by unpaired two tail t-test.

### Effects of fosfomycin resistance mutations on the transcriptional profile of *S. maltophilia*

In order to know if the mutation of genes encoding the enzymes of the central metabolism change the transcriptional profile in a way directly related to fosfomycin resistance, the transcriptomes of the fosfomycin resistant mutants were compared to that of the wild-type strain. Changes in the expression levels of just 67 of the 4210 genes that form the genome of *S. maltophilia* D457 were detected (Table S2). Most changes were specific for each mutant, indicating that the observed transcriptomic changes were unlikely associated to the common phenotype of fosfomycin resistance (Figure S4). Concerning changes that may explain the resistance phenotype is important to note the absence of relevant transcriptional changes in genes related to cell wall synthesis, such as the gene encoding the fosfomycin target MurA and SMD_1053, SMD_1054, SMD_0334, *nagZ* and SMD_2885, predicted to be involved in the recycling of the peptidoglycan (Table S3). These results support that an increased expression of either the fosfomycin target (MurA) or the alternative peptidoglycan recycling pathway is not the cause of fosfomycin resistance in the analyzed mutants.

## Discussion

So far described fosfomycin resistance mechanisms can be clustered into three classical categories of antibiotic resistance acquisition (1): alterations in fosfomycin transport, antibiotic inactivation and alterations in the target enzyme or peptidoglycan biosynthesis (17). Herein, using a set of *in vitro* selected mutants, we have shown that none of these already known mechanisms seem to be involved in the acquisition of mutation-driven fosfomycin resistance by *S. maltophilia*. In this microorganism, the acquisition of resistance is due to the inactivation of enzymes belonging to the EMP pathway.

Our results indicate that the inactivation of these enzymes does not cause major changes in the transcriptomes of the mutants that may justify resistance as the consequence of a collateral effect of the selected mutations on the expression of the aforementioned fosfomycin resistance mechanisms. Inasmuch, intracellular accumulation of fosfomycin was similar in the wild-type and the mutant strains, which support that resistance is neither due to an impaired fosfomycin uptake (25) nor to its degradation *via* the activity of fosfomycin-inactivating enzymes (29, 53). Also supporting this result is the fact that the genome of *S. maltophilia* does not encode homologous of the already known fosfomycin resistance proteins or its transporters GlpT and UhpT.

Other mechanisms leading to fosfomycin resistance are modifications of the target MurA (21) or changes in its expression level. Nevertheless, when the mutants were sequenced, no mutations in *murA* were found and the analysis of the transcriptomes indicates that *murA* is not expressed at higher levels in the resistant mutants than in the wild-type strain. Same happens with the pathway involved in the recycling of peptidoglycan, which increased expression may contribute to fosfomycin resistance (7). The expression of the genes encoding the enzymes of this pathway is not higher in the mutants that in the wild-type strains as shown in the transcriptomic studies.

Therefore, classical antibiotic resistance mechanisms (1, 54) do not seem to be the cause of fosfomycin resistance in *S. maltophilia*. Although, at above stated, there are not relevant transcriptional changes in the mutants, these strains appear to show a different physiological state than the wild-type strain, as evidenced by the fact that they exhibit increased Zwf activity together with the loss of function of the mutated enzymes. These changes do not modify the response to oxidative stress, an element that could be relevant in the activity of antibiotics (39). However, it is worth mentioning that the regulation of the metabolic fluxes of carbon metabolism include additional mechanisms other than transcriptional regulation (55). Among them, allosteric regulation as well as the activity of posttranscriptional or posttranslational regulators can change the production levels and activity of different proteins (eventually involved in the resistance phenotype) without changing their mRNA levels (56).

The mutated enzymes belong to the amphibolic metabolic pathway (EMP and gluconeogenesis) which includes PEP, the natural substrate of MurA. Fosfomycin due to its structural similarities to PEP binds and inhibits MurA. It might be then possible that inactivation of such enzymes in the fosfomycin resistant mutants may produce an increased synthesis of PEP that could outcompete fosfomycin for its binding to MurA. However, the concentrations of PEP are no higher in the fosfomycin resistant mutants than in the wild-type strain, which goes against this possibility. Several enzymes from central metabolism are moonlighting proteins; they display functions unrelated to their enzymatic activity (42). The complementation of the mutants with *E. coli* enzymes restored their susceptibility to fosfomycin, which indicates that the impaired activity of these metabolic enzymes is on the basis of the observed phenotype of fosfomycin resistance. Nevertheless, a possible function of the *S. maltophilia* enzymes not related to their known metabolic function cannot be totally discarded.

Previous analysis has shown that *E. coli* mutants deficient in the metabolic enzyme isocitrate dehydrogenase are resistant to nalidixic acid (35). However, little work is still available in the crosstalk between metabolism (and metabolic robustness) and antibiotic resistance (57, 58), despite the fact that metabolic interventions may improve the activity of the antibiotics (33, 59-61) and that bacterial metabolism can constrain the evolution of antibiotic resistance (13). Our results highlight the importance that the modification of the activity of enzymes belonging to central metabolism may have in the susceptibility to antibiotics, as fosfomycin, that are not known to interact with such enzymes. The finding that fosfomycin activity is highly dependent on the bacterial metabolic status, being more active when bacteria grow intracellularly (27, 28) or under acidic conditions and anaerobiosis in urine (62), further support that antibiotic activity and, consequently antibiotic resistance, are interlinked with the bacterial metabolism.

## Material and methods

### Bacterial strains and culture conditions

All bacterial strains, plasmids and oligonucleotides and used in this study are listed in Tables S4 and S5. Unless otherwise stated, bacteria were grown in LB (Lysogeny Broth) Lennox medium at 37 °C with constant agitation at 250 rpm. Solid medium was prepared using an agar concentration of 15 g/l. In order to analyze the growth of *S. maltophilia* D457 in the presence of a single carbon source, *S. maltophilia* minimum medium (SMMM) (63) with some modifications was used. The composition of the medium is described in Supplemental Materials and Methods. When required, antibiotics were added: 100 μg/ml ampicillin and 50 μg/ml kanamycin for *E. coli*, and 500 μg/ml kanamycin for *S. maltophilia*. Besides, different concentrations of fosfomycin as well as 1 mM of IPTG were used in different experiments, as stated in the different sections.

### Isolation of fosfomycin resistant mutants

Around 10^8^ *S. maltophilia* D457 bacteria cells were plated on MH agar Petri dishes containing 1024 μg/ml fosfomycin and were grown at 37 °C during 48 h. The mutants selected in these conditions were grown on LB agar without antibiotic (three sequential passages) and then were grown again on MH agar containing 1024 μg/ml fosfomycin to ensure that the observed phenotype was not transient. The susceptibility of mutants to fosfomycin was tested (see below), for further studies 4 mutants were randomly selected and dubbed FOS1, FOS4, FOS7 and FOS8.

### DNA extraction, whole genome sequencing and SNP identification

Chromosomal DNA from each mutant (FOS1, FOS4, FOS7 and FOS8) and the wild-type strain (D457) was obtained from overnight cultures using the GNOME DNA kit (MP Biomedicals). Genomic DNAs were sequenced using the Illumina technology at the Parque Científico of Madrid, Spain. The samples were subjected to single-end sequencing with a read-length of 1×150 and a coverage between 26 and 41X was obtained. The genomic sequences of the strains were compared with *S. maltophilia* D457 reference genome (NC_017671.1) and visualized using the software FIESTA 1.1 (http://bioinfogp.cnb.csic.es/tools/FIESTA). Mutations were filtered according to sequence quality (>30) and the mutation effect in the protein sequence (moderate and high effect), and the variants absent in the control D457 parental strain were studied. Provean predictor (provean.jcvi.org) was used to anticipate whether an amino acid substitution or indel had an impact on the biological function of the coding protein.

The presence of the mutations detected from the whole genome sequencing analysis was confirmed as described in Supplemental Materials and Methods.

### Antimicrobial susceptibility assays

The minimal inhibitory concentrations (MICs) of gentamicin, tobramycin, ciprofloxacin, nalidixic acid, ceftazidime, colistin, tetracycline, chloramphenicol and fosfomycin were determined for each strain on LB agar using MIC test strips (MIC Test Strips, Liofilchem Diagnostics). For phenotypic the analysis of mutants complemented with the Glucobrick module II, antibiotic disks (Oxoid) were used. Plates were incubated at 37 °C and results were analyzed after 20 h. Since commercial fosfomycin disks contain glucose 6-P, fosfomycin susceptibility assays were also performed, under the same growing conditions, using paper disks (9 mm, Machery-Nagel) impregnated with 0.5 mg of fosfomycin. The experiments were performed in triplicate.

### Complementation of fosfomycin resistant mutants and generation of *zwf* deletion mutants

The genes *eno, gpmA, pgk* and *gapA*, encoding glycolytic enzymes, were obtained from the wild-type strain *S. maltophilia* D457 by PCR amplification and introduced in *S. maltophilia* as described in Supplemental Material and Methods.

To complement the mutants with a partial version of the Glucobrick module II, which contains the genes of the lower glycolysis enzymes of *E. coli* K12, (*gapA, pgk, gpmA, eno* and *pyk*) (64), the pSEVA224 GBII plasmid containing these genes was purified with the QIAprep Spin Miniprep kit and digested with restriction enzymes BamHI and HindIII, obtaining the *gapA-pgk-gpmA-eno* fragment of the Glucobrick module II. The corresponding band was purified and ligated into pSEVA234 previously digested with the same enzymes. The new pSEVA234(*gapA-pgk-gpmA-eno*) plasmid was introduced into the *S. maltophilia* strains D457, FOS1, FOS4, FOS7 and FOS8 by triple conjugation (65) as described in Supplemental Materials and Methods.

The *zwf* gene was deleted in different *S. maltophilia* strains by homologous recombination as described (66) and detailed in the Supplemental Materials and Methods section.

### RNA extraction and RNA-Seq

The different bacterial strains were grown overnight in LB broth at 37 °C and 250 rpm. These cultures were used to inoculate new flasks to reach an 0.01 OD_600_ and the cultures were grown at 37 °C until an OD_600_ of 0.6 was reached. Afterwards, RNA was isolated (67) and RNA-Seq was carried out at Sistemas Genómicos S.L., Parque Tecnológico de Valencia, with Illumina HiSeq 2500 sequencing technology using a 50 bp single-end format as described in Supplemental Materials and Methods.

### Bacterial growth measurements

Growth was measured with a *Spark 10M* plate reader (TECAN) at OD_600_ in flatbottomed transparent 96-well plates (Nunc MicroWell Thermo Fisher). Each well was inoculated with bacterial suspensions to a final OD_600_ of 0.01 in SMMM containing 40 mM of the carbon source under study or LB medium. For SMMM experiments, overnight cultures were washed twice with SMMM medium without any carbon source. The plates were incubated at 37 °C with 10 s of shaking every 10 min.

### Protein quantification

Protein concentration was determined following the Pierce BCA Protein Assay Kit (Thermo Scientific) protocol in 96 well plates (Nunc MicroWell Thermo Fisher).

### *In vitro* activity assays of the lower glycolysis enzymes and dehydrogenases

Cells were harvested at exponential phase (OD_600_ = 0.6) by centrifugation at 5100 x*g* and 4 °C and washed twice in 0.9% NaCl and 10 mM MgSO_4_. Once washed, cells were disrupted by sonication at 4 °C and the cell extracts were obtained by centrifugation at 23100 x*g* for 30 min at 4 °C.

NAD(P)^+^ reduction or NAD(P)H oxidation was monitored spectrophotometrically at 340 nm and 25 °C with intermittent shaking in microtiter plates using *Spark 10M* plate reader (TECAN). Each reaction was performed using three biological replicates and the specific activities were obtained by dividing the measured slope of NAD(P)H formation or consumption by the total protein concentration. Enzymatic activities of dehydrogenases (glucose-6-phosphate, isocitrate and glyceraldehyde-3P dehydrogenases) were measured as described (68). Enzymatic activities of phosphoglycerate kinase, phosphoglycerate mutase and enolase were assayed following the protocol described by Pawluk, A. *et al* (69) with some modifications in a two-step reaction (see Supplemental Materials and Methods).

### Quantification of intracellular phosphoenolpyruvate and fosfomycin

The amount of PEP was measured from cultures in exponential growth phase in LB medium (OD_600_ = 0.6). Twenty ml of each culture were centrifuged at 4500 x*g* for 3 min at 4 °C. *PEP Colorimetric / Fluorometric Assay Kit* protocol (Sigma-Aldrich) was used with some modifications that are described in Supplemental Materials and Methods.

Assays to test fosfomycin accumulation in bacterial cells were conducted as previously stated (49), with some modifications that are described in Supplemental Materials and Methods.

### H_2_O_2_ and menadione susceptibility test

The susceptibility to H_2_O_2_ and menadione was tested as described previously with some modifications (70). Sterile paper disks (9 mm, Machery-Nagel) were impregnated with 10 μl of 2.5% H_2_O_2_ or 20 μl of 0.2 M menadione and placed on LB agar plates. The diameter of the zone of growth inhibition around each disk was measured after 20 h of incubation at 37 °C. The experiment was performed in triplicate.

### Metabolic map of *S. maltophilia*

To model the metabolic map of *S. maltophilia* D457, indicating possible enzymes of the central metabolism and route bypasses, the BioCyc database (71) was used. The sequence of the enzymes was obtained from the complete genome of *S. maltophilia* D457 (72). In addition, the amino acid sequence of the enzymes of the central metabolism of *P. aeruginosa* PAO1 (73) and *E. coli* (74) were aligned using the Blast tool (48) with the *S. maltophilia* D457 genome confirming the presence or absence of these enzymes. Moreover, Blast was used to identify possible peptidoglycan recycling pathway genes, fosfomycin transporters and fosfomycin resistance proteins in *S. maltophilia* D457.

## Acknowledgements

Work in the laboratory is supported by Instituto de Salud Carlos III (grant RD16/0016/0011) - cofinanced by the European Development Regional Fund “A Way to Achieve Europe”, by grant S2017/BMD-3691 InGEMICS-CM, funded by Comunidad de Madrid (Spain) and European Structural and Investment Funds and by the Spanish Ministry of Economy and Competitivity (BIO2017-83128-R). TGG is the recipient of a FPI fellowship from MINECO. The funders had no role in study design, data collection and interpretation, or the decision to submit the work for publication.

